# Topographic axonal projection at single-cell precision supports local retinotopy in the mouse superior colliculus

**DOI:** 10.1101/2022.03.25.485790

**Authors:** Dmitry Molotkov, Leiron Ferrarese, Tom Boissonnet, Hiroki Asari

## Abstract

Retinotopy, like all long-range projections, can arise from the axons themselves or their targets. The underlying connectivity pattern, however, remains elusive at the fine scale in the mammalian brain. To address this question, we functionally mapped the spatial organization of the input axons and target neurons in the mouse retinocollicular pathway at single-cell resolution using *in vivo* two-photon calcium imaging. We found a near-perfect retinotopic tiling of retinal ganglion cell axon terminals, with an average error below 30 μm or 2 degrees of visual angle. The precision of retinotopy was relatively lower for local neurons in the superior colliculus. Subsequent data-driven modelling ascribed it to a low input convergence, on average 5.5 retinal ganglion cell inputs to a postsynaptic cell in the superior colliculus. These results indicate that retinotopy arises largely from topographically precise input from presynaptic cells, rather than elaborating local circuitry to reconstruct the topography by postsynaptic cells.

## Introduction

Topographic organization is central to brain function (Jbabdi et al., 2013; Patel et al., 2014). Connections between brain regions are often spatially arranged to have one-to-one mappings, and sensory processing relies on the transmission of topographically preserved information from receptor organs. In the visual system, for example, topographic visual representations are first formed in the retina and conveyed to the brain by retinal ganglion cells (RGCs) whose axons are bundled into an optic nerve (Masland, 2012). In general, neighboring RGCs project to neighboring regions in their targets (Frisén et al., 1998; McLaughlin et al., 2003; Cang and Feldheim, 2013; Arroyo and Feller, 2016). This forms a retinotopic map in the primary retinorecipient areas, such as the lateral geniculate nucleus (Malpeli and Baker, 1975) and the superior colliculus (SC; Dräger and Hubel, 1976; Cang et al., 2018), and retinotopy is likewise transferred throughout the entire visual pathways (Wandell et al., 2007; Cang and Feldheim, 2013).

How precisely is topographic information transmitted between brain regions? Despite the substantial progress in our understanding of the connectivity patterns in simple nervous systems (Varshney, et al., 2011; Kunst, et al., 2019; Winding, et al., 2023), only global-level relationship is known even for the best characterized cases in mammals, such as the retinal projection to SC (Frisén et al., 1998; McLaughlin et al., 2003; Cang and Feldheim, 2013; Arroyo and Feller, 2016). During development, RGC axons reach their target locations based on molecular guidance cues and activity-dependent refinement (Chandrasekaran et al 2005; Plas et al., 2005; Dhande et al., 2011). Long-range projections of dense axonal fibers, however, preclude a precise anatomical characterization of the connectivity patterns at single-cell resolution (Hong et al., 2011). In the optic nerve, RGC axons are mixed and lose retinotopic organizations (Horton et al., 1979; Colello and Guillery, 1998). While pretarget sorting of the axons partially restores the topography in the optic tract (Plas et al., 2005; Cioni et al., 2018), a fundamental question is then if RGC axons can nevertheless find precise targets even after they get “lost” in the long-range projection (Figure 1, Model 1), or if their target neurons need to reconstruct the topography by elaborating local circuitry (Figure 1, Model 2). A recent study using a high-density electrode has implicated retinotopic RGC projections to the mouse SC along the probe shank (Sibille et al., 2022), favoring the former scenario. However, the precision of retinotopy at the level of individual axons and its relationship to that of target neurons remain elusive in any retinorecipient area of the mammalian nervous system.

**Figure 1:**
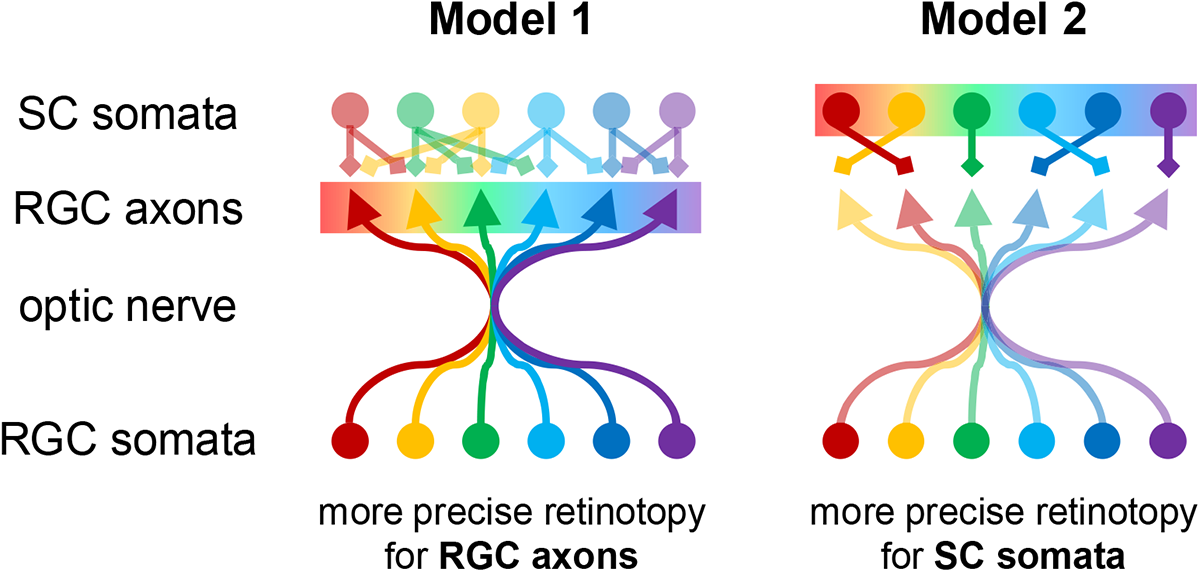
Possible retinocollicular projection models for topographic information transmission. Retinotopy in SC (color-coded) can arise from either precisely retinotopic projection of RGC axons, innervating exact target locations in SC (Model 1); or roughly 49 retinotopic projection of RGC axons while SC neurons identify appropriate partners to recover the retinotopy (Model 2). In Model 1, retinotopy can be more precise for RGC axons than for SC somata if the synaptic connectivity is not topographically well organized. In Model 2, in contrast, SC somata should show more precise retinotopy than RGC axons.

To address this question, we performed functional mapping of RGC axon terminals and local neurons in SC at single-cell resolution in awake head-fixed mice using two-photon calcium imaging. Subsequent analyses on their spatial organizations allowed us to quantify and directly compare the precision of retinotopy between the pre- and post-synaptic sides. Moreover, these experimental data served as a basis of a computational modelling analysis for inferring key features on the topographic information transmission in the retinocollicular pathway. Specifically, by comparing the observed and simulated retinotopy patterns, here we addressed 1) to what extent RGC axon terminals are deviated from their retinotopically optimal target locations in SC; and 2) on average how many RGCs connect to individual SC neurons.

## Results

For functional mapping of the mouse retinocollicular projections, we expressed axon-targeted calcium indicators (GCaMP6s; Broussard et al., 2018) in RGCs via intravitreal injection of recombinant adeno-associated viruses (AAVs) harboring the pan-neuronal human synapsin (hSyn) promoter, and monitored the visual responses of the labelled RGC axons in SC using *in vivo* two-photon microscopy (Figure 2A,B and Supplemental Movie 1). To segment axonal patches of individual RGCs and isolate their activity, we used constrained non-negative matrix factorization (CNMF) that allows us to extract morphological and temporal features from noisy time-lapse calcium activities based on their spatiotemporal correlations (Giovannucci et al., 2019). For a field of view of ∼0.3 mm^2^ (0.57-by-0.57 mm), we detected 26 ± 9 axonal patches displaying independent activity patterns (median ± median absolute deviation here and thereafter unless otherwise noted; N=37 recording sites in total from 15 animals; e.g., Figure 2C,D). The size of the individual axonal patches (135 ± 25 μm; N=969; Supplemental Figure 1A) is consistent with the past anatomical measurement (Hong et al., 2011), with a substantial overlap between them over the SC surface (49 ± 11 %). This supports a successful signal extraction and a good coverage of RGC axonal labelling. These data allowed us to faithfully reconstruct the local two-dimensional (2D) map of the individual RGC axon terminals in SC (e.g., Figure 2D).

**Figure 2:**
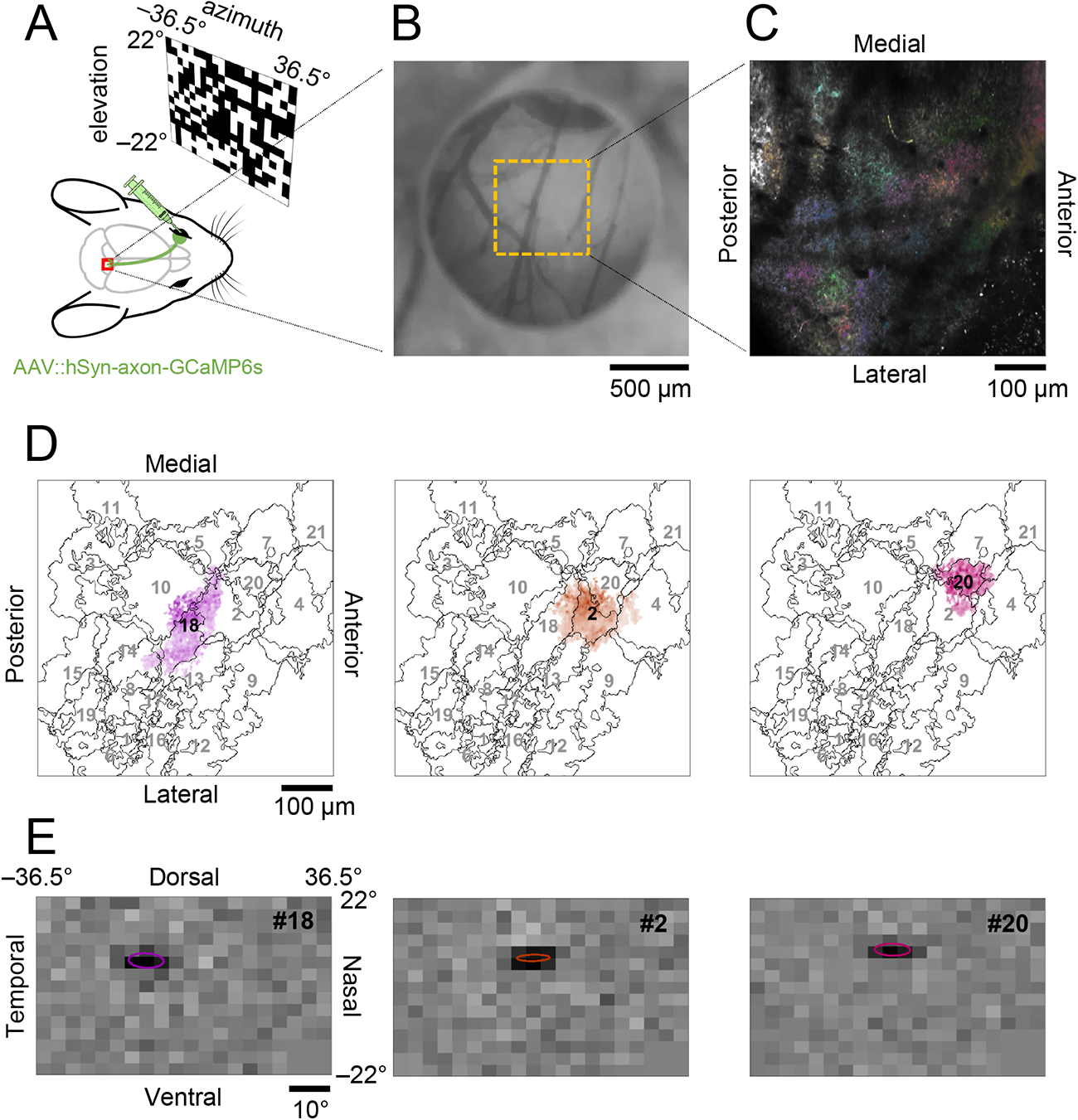
*In vivo* two-photon calcium imaging of retinal ganglion cell axon terminals in the mouse superior colliculus. *A*: Schematic diagram of the experimental set up for RGC axonal imaging in SC. *B*: Representative image of cranial window. Medial-posterior part of SC was clearly visible through a cylindrical silicone plug attached to a glass coverslip. *C*: Average intensity projection of representative axonal imaging data, overlaid with detected RGC axonal patches (N=21; color-coded). See also Supplemental Movie 1. *D*: Footprint of three representative RGC axonal patches (#18, 2, and 20 in distinct color; from left to right, respectively) overlaid with the profile of the rest patches (in grey). *E*: Corresponding RF of the three representative RGC axonal patches (from D), estimated by reverse-correlation analysis (ellipse, 1 SD Gaussian profile).

**Supplemental Figure 1:**
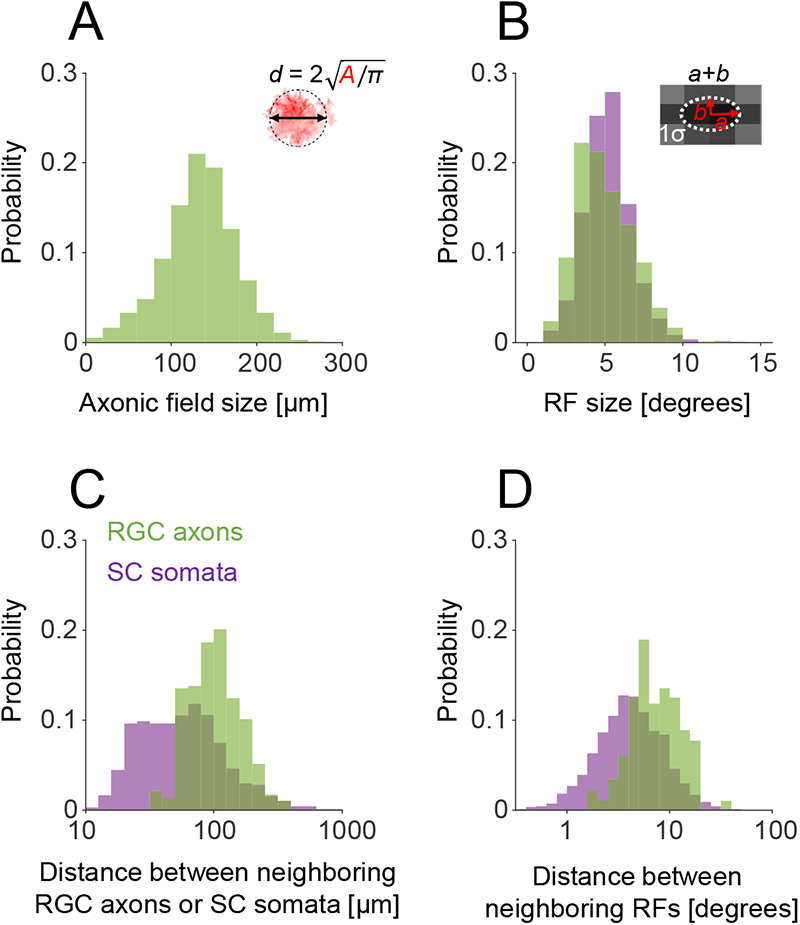
Probability distributions of axonic and receptive field statistics. *A*: Size of the identified RGC axon terminals, d = 2(A/π)0.5, where A is the axonic field area from the CNMF analysis (135 ± 25 μm, median ± median absolute deviation; N=969). *B*: RF size of RGC axons (green; 4.8 ± 1.2 degrees; N=719) and SC somata (purple; 5.1 ± 0.9 degrees; N=1191), measured as the mean of long- and short-diameters (equivalently, the sum of long- and short-radii, a and b, respectively) of the 2D Gaussian profile at 1 σ fitted to the RF (e.g., Figure 2E). *C*: Distance between neighboring RGC axon centers (green; 100 ± 30 μm; N=761 pairs) or SC somata (purple; 56 ± 28 μm; N=2292 pairs). Neighboring pairs are identified by the Delaunay triangulation analysis (e.g., Figure 3C and Supplemental Figure 3F). *D*: Distance between neighboring RF centers of RGC axons (green; 7.2 ± 2.7 degrees; N=776 pairs) or SC somata (purple; 4.1 ± 1.9 degrees; N=2287 pairs). Neighboring pairs are identified by the Delaunay triangulation analysis (e.g., Figure 3D and Supplemental Figure 3G).

**Supplemental Movie 1:**
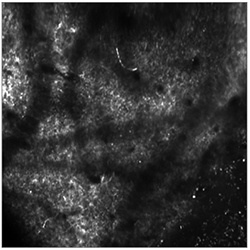
*In vivo* two-photon calcium imaging of retinal ganglion cell axon terminals in the mouse superior colliculus. A snippet of rigid motion-corrected movie showing the activity of RGC axons in the mouse SC in response to the random checkerboard stimuli (sampling rate, 15.4 Hz; averaged every 5 frames for noise reduction, hence sped up by 5 times). Remaining non-rigid motion was further corrected before signal extraction by CaImAn (see Methods for details). A black square overlaid at the bottom-right corner turns white when the visual stimulus is on. See Figure 2 for the segmentation and Figure 3 for the tiling pattern analysis.

Presence of a well-defined receptive field (RF) is a characteristic feature of all RGC types (Masland, 2012; Baden et al., 2016). To map the RF of the identified RGC axons, we computed the response-weighted average of the presented random checkerboard stimuli (frame rate, 4 Hz; rectangular fields, 3.7° in width and 2.9° in height; e.g., Figure 2E) and fitted a 2D Gaussian at the peak latency to characterize the spatial RF profile. Most identified axonal patches had RFs within the stimulation screen (±22° in elevation and ±36.5° in azimuth from the mouse eye; Boissonnet et al., 2022). In accordance with *ex vivo* retinal physiology (Baden et al., 2016), the average RF size of the RGC axons was 4.8 ± 1.2 degrees (N=719; Supplemental Figure 1B), estimated as the mean of the long- and short-axis diameters of the 2D Gaussian profile at 1 standard deviation (SD). There was a weak but statistically significant correlation between the RF size and the RGC axonal patch size (Pearson’s r = 0.19, p = 5e-7). The RFs locally tiled the visual field with 10 ± 5 % overlap at 1 SD Gaussian profiles, where the center of every RF occupied a unique location in the visual field. This ensures that these RFs belong to different RGCs because the RF center location of RGC axons should correspond well to the location of their somata in the retina. Hence, the RF tiling faithfully represents retinotopy.

How well does the tiling pattern of RGC axons in SC agree with that of their RFs? As expected from the global retinotopy in SC (Dräger and Hubel, 1976; Cang et al., 2018; Sibille et al., 2022), relative positions of the RGC axonal patches (e.g., Figure 3A) agreed well with those of the corresponding RFs (e.g., Figure 3B) regardless of their cell types. For quantification, we first computed the Delaunay triangulation using the geometric centers of the individual axonal patches or the RF center locations as landmark points in each space (e.g., Figure 3C,D, respectively). This triangulation features the adjacency relationship regardless of their absolute positions, where all the adjacent pairs of the landmark points in a given space are connected as a dual graph of the Voronoi tessellation that separates the space into territories close to each landmark point. The distances between the centers of neighboring RGC axons and those between their RFs were identified to be 100 ± 30 µm (N=761 pairs; Supplemental Figure 1C) and 7.2 ± 2.7 degrees (N=776 pairs; Supplemental Figure 1D), respectively. As a measure of the agreement between the two tiling patterns, we then calculated the fraction of the common edges between the two Delaunay triangulations (blue edges; 88.2% for the example in Figure 3C,D). Throughout our datasets, we found a near-perfect match between the tiling patterns of RGC axons in SC and their RFs (84 ± 5%; N=36 recordings from 12 animals; Figure 4A) regardless of the recording depth within the superficial SC layer (120-220 μm deep from the surface; Figure 4B). This observation is consistent with the precise axonal projection model (Model 1 in Figure 1) whereby RGC axon terminals retinotopically tile the SC surface at single-cell precision.

**Figure 3:**
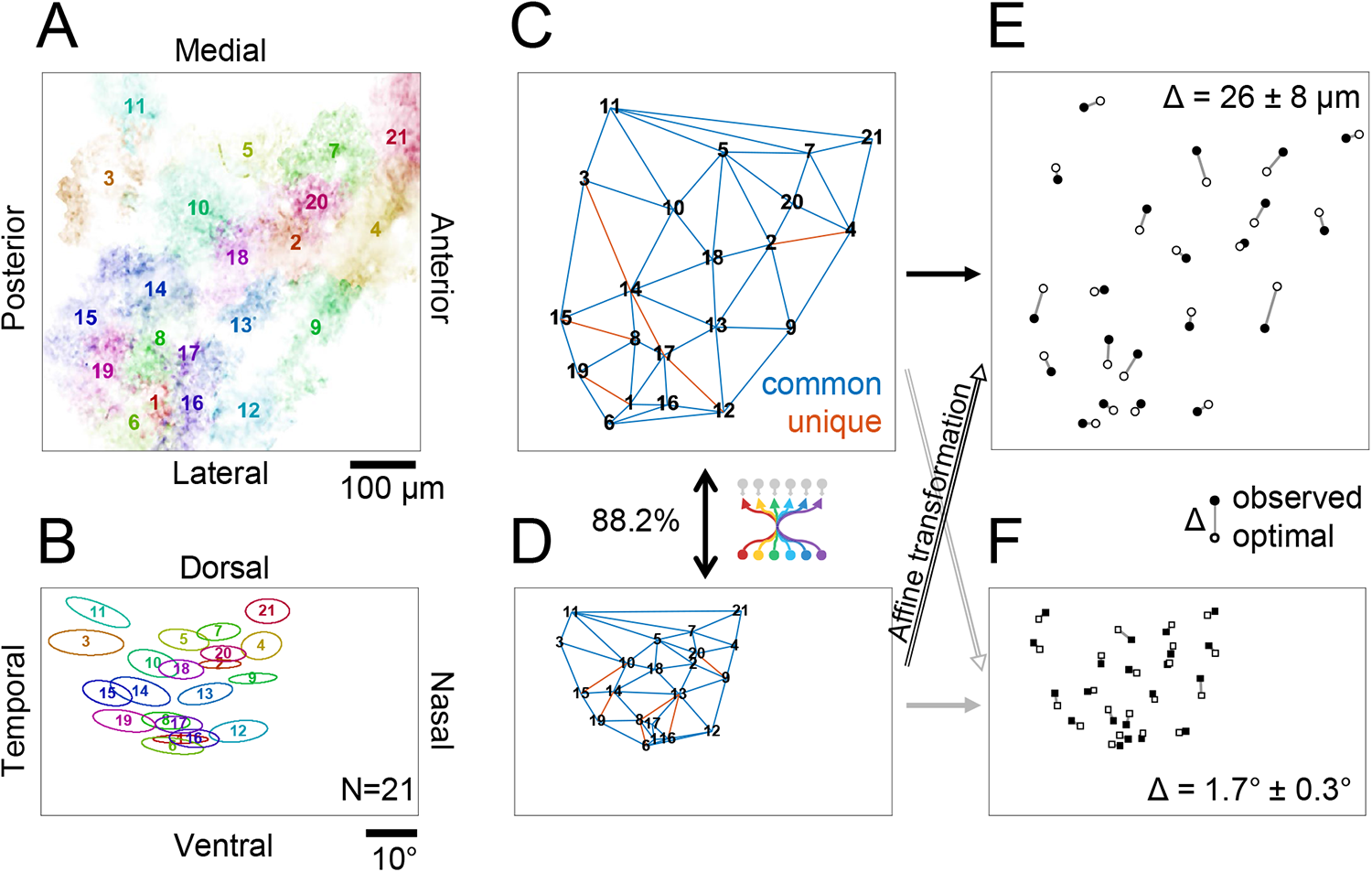
Retinal ganglion cell axons retinotopically tile the mouse superior colliculus at single-cell precision. See Supplemental Figure 2 for another example. *A,B*: RGC axon patches in SC from a representative recording session (A; N=21, color-coded; from Figure 2C) and their corresponding RFs (B; 1 SD Gaussian profile). *C,D*: Delaunay triangulation of the RGC axonal patch locations (C; from A) and the RF centers (D; from B). The two triangulation patterns are nearly identical (88.2%; blue, common edges in both patterns; red, unique edges only in either pattern). *E*: Comparison between the observed tiling pattern of RGC axons (filled circles; from C) and the retinotopically ideal pattern (open circle) obtained by applying an optimal Affine transformation to the corresponding RF locations (in D) that minimizes the discrepancy between the two patterns (Δ = 26 ± 8 μm). *F*: Corresponding comparison between the observed (filled squares; from D) and ideal (Affine-transformed pattern in C) RF tiling patterns of RGC axons (Δ = 1.7 ± 0.3 degrees).

**Supplemental Figure 2:**
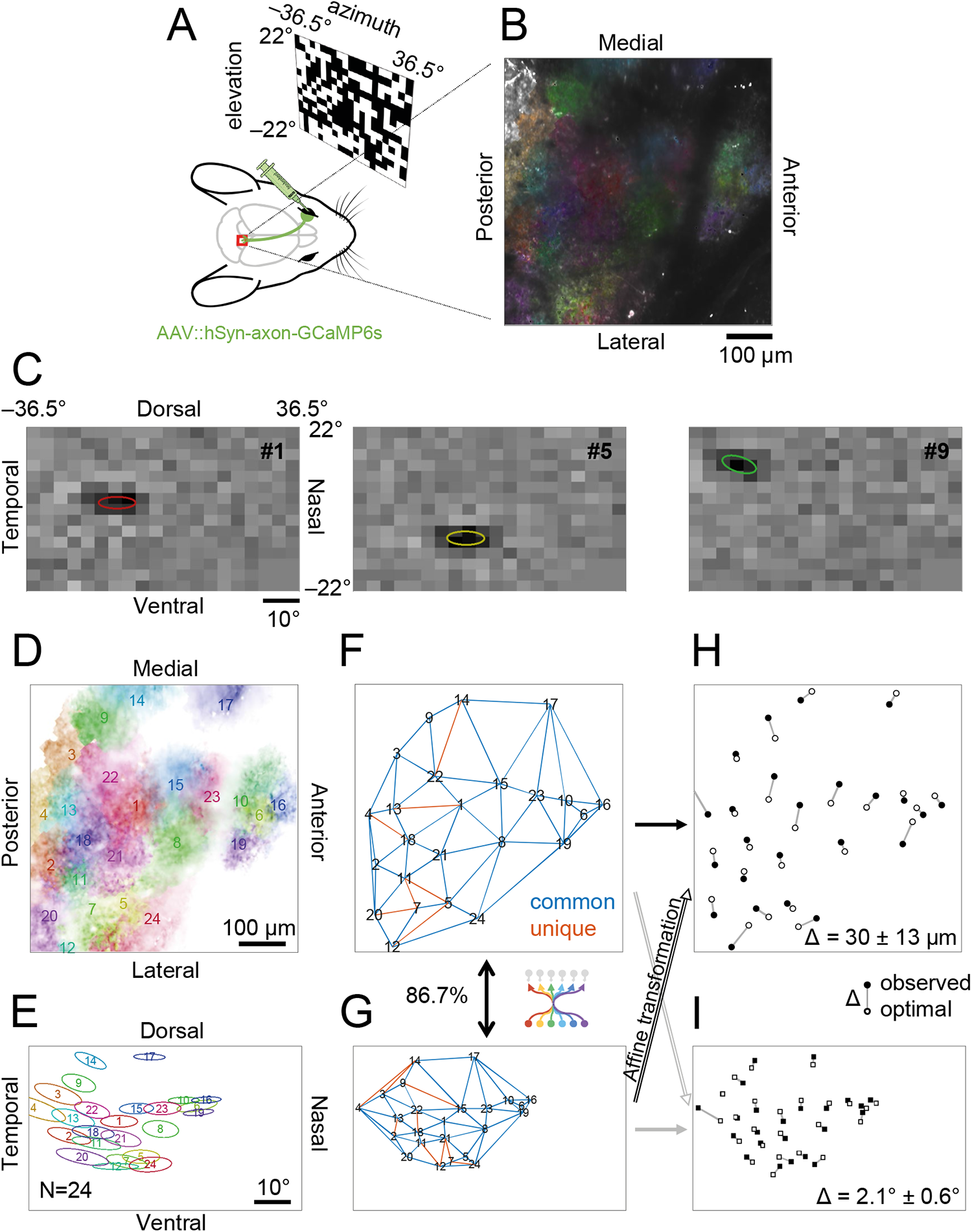
Another example of precise tiling of retinal ganglion cell axons in the mouse superior colliculus. Figure panels are shown in the same format as in Figures 2 and 3. *A*: Schematic diagram of RGC axonal imaging in SC. *B*: Average intensity projection of representative imaging data, overlaid with detected RGC axonal patches (N=24; color-coded). *C*: RFs of three representative RGC axonal patches (ellipse, 1 SD Gaussian profile). *D,E*: Tiling patterns of RGC axons (D) and their corresponding RFs (E; 1 SD Gaussian profiles with the same color-code as in B). *F,G*: Delaunay triangulation of the RGC axon centers (F; from D) and the RF centers (G; from E), showing a good agreement between them (86.7%; blue, common edges in both patterns; red, unique edges only in either pattern). *H,I*: Comparison between the observed and retinotopically ideal tiling patterns of RGC axons (H; Δ = 30 ± 13 μm) or their RFs (I; Δ = 2.1 ± 0.6 degrees).

**Supplemental Figure 3:**
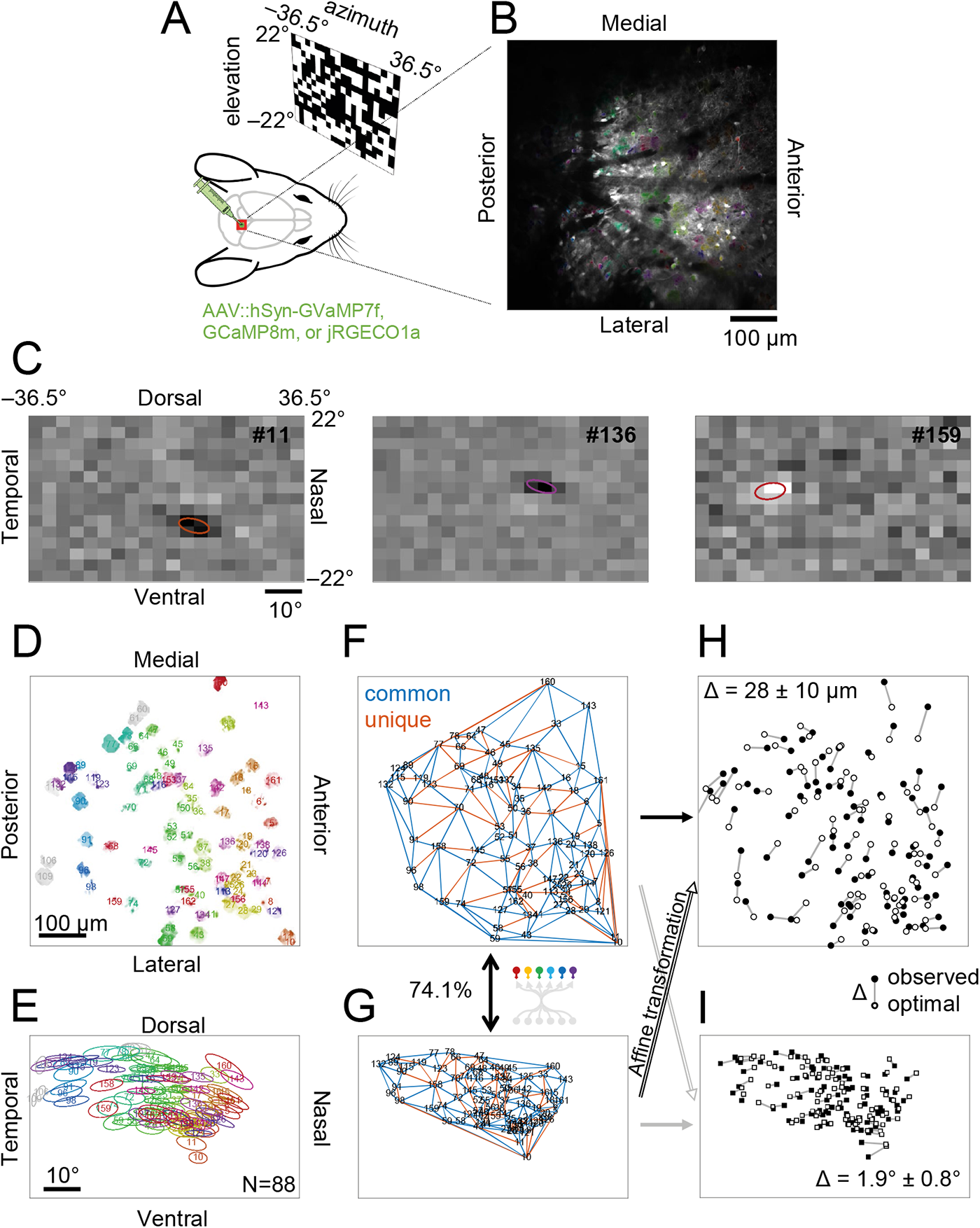
Tiling pattern analysis of local neurons in the mouse superior colliculus. Figure panels are shown in the same format as in Figures 2 and 3. *A*: Schematic diagram of the experimental set up for SC somatic imaging. *B*:Average intensity projection of representative imaging data, overlaid with detected SC somata (N=88; color-coded). *C*: Representative RFs of local SC neurons (from left to right: #11, #136, #159) from reverse-correlation analysis (ellipse, 1 SD Gaussian profile). *D,E*: Tiling pattern of SC somata (D) from a representative recording session (B) and their corresponding RF tiling pattern (E; 1 SD Gaussian profiles with the same color-code as in B). *F,G*: Delaunay triangulation of the SC somatic locations (F; from D) and the RF centers (G; from E), showing a good agreement between them (74.1%; blue, common edges in both patterns; red, unique edges only in either pattern). *H,I*: Comparison between the observed and retinotopically ideal tiling patterns of SC somata (H; Δ = 28 ± 10 μm) or their RFs (I; Δ = 1.9 ± 0.8 degrees).

**Figure 4:**
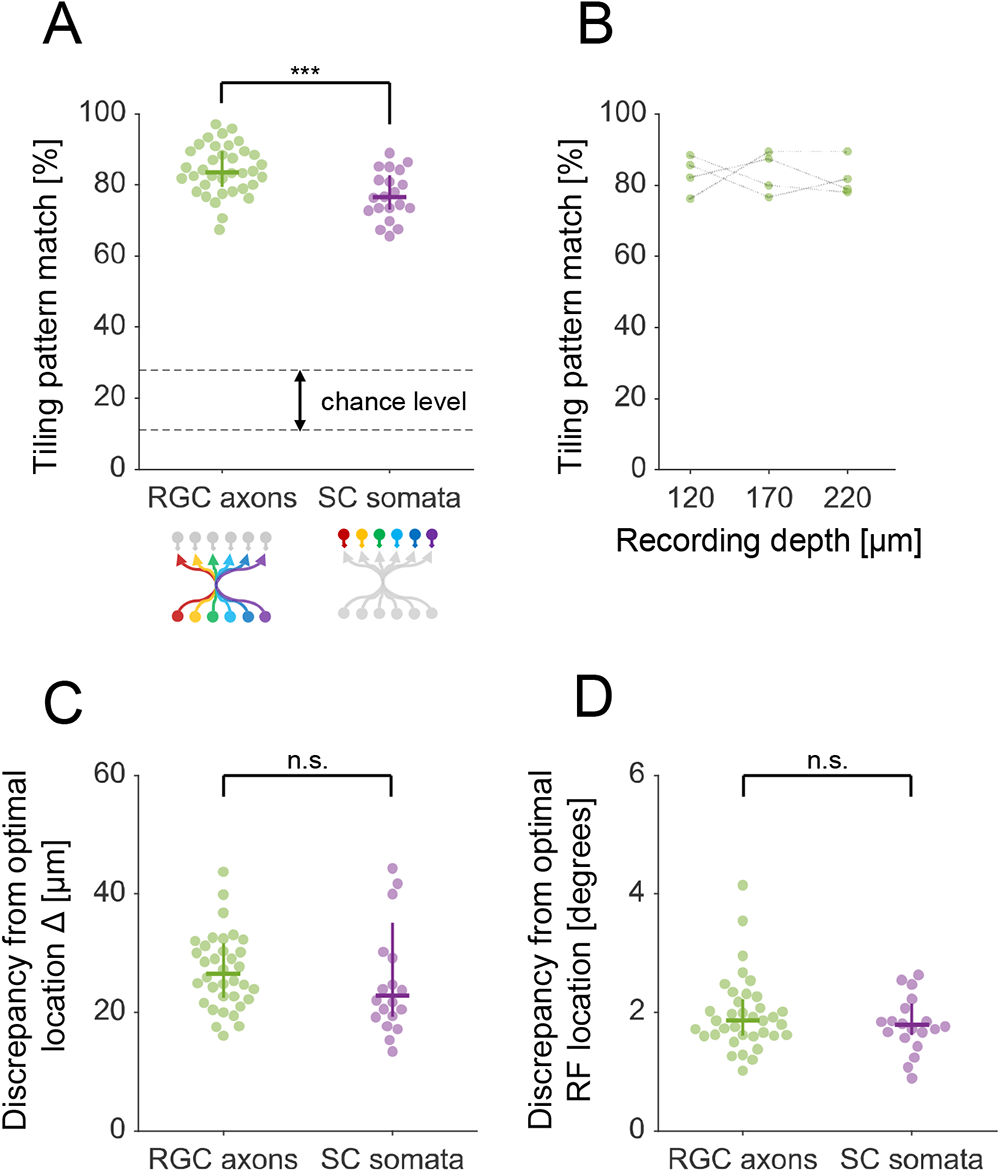
Tiling patterns of retinal ganglion cell axons are retinotopically more precise than those of local neurons in the mouse superior colliculus. *A*: RGC axons (84 ± 5%; N=36 recordings from 12 animals; e.g., Figure 3) showed a significantly higher precision of retinotopy than SC somata (77 ± 5%; N=20 recordings from 20 animals; e.g., Supplemental Figure 3); p = 0.001, rank sum test. The error bars show the median and interquartile range; and the dotted lines represent the chance level at p = 0.05 (11-28%; bootstrap with 10,000 repetitions). *B*: The precision of the retinotopic tiling of RGC axons was not dependent on the projection depth from the SC surface (N=4 animals; 83 ± 5, 83 ± 6, and 82 ± 5% at 120, 170, and 220 μm deep, respectively; mean ± SD; p = 0.9, one-way analysis-of-variance). *C*: Deviation between the observed and retinotopically ideal locations of the RGC axons (27 ± 4 μm; e.g., Figure 3E) or SC somata (23 ± 5 μm; e.g., Supplemental Figure 3H); p = 0.27, rank sum test. *D*: Deviation between the observed and retinotopically ideal RF locations of the RGC axons (1.9 ± 0.3 degrees; e.g., Figure 3F) or SC somata (1.8 ± 0.3 degrees; e.g., Supplemental Figure 3I); p = 0.66, rank sum test.

To further quantify the precision of the RGC axonal projection, we compared the observed position with the ideal one that forms perfect retinotopy (e.g., Figure 3E). Specifically, we used a linear method to estimate this retinotopically optimal tiling pattern of RGC axons: i.e., an Affine transformation that best mapped the observed RF tiling pattern onto the corresponding observed axonal tiling pattern (see Methods for details). We found that the average discrepancy between the observed and retinotopically optimal RGC axonal locations in SC was 27 ± 4 μm (Figure 4C). This is much shorter than the distance between neighboring RGC axon centers (100 ± 30 μm, N=761 pairs; Supplemental Figure 1C) or the axonal patch size (135 ± 25 μm, N=969; Supplemental Figure 1A), and thus will not have a measurable impact on the retinotopy. Likewise, the extent to which the observed RF tiling pattern deviated from the linear optimal one (1.9 ± 0.3 degrees; Figure 4D; see Figure 3F for example) was much smaller than the RF size of RGC axons (4.8 ± 1.2 degrees, N=719; Supplemental Figure 1B) or the spacing between the RFs (7.2 ± 2.7 degrees, N=776 pairs; Supplemental Figure 1D). These data support that RGCs can precisely innervate their axons to their target locations and faithfully transmit the information about retinotopy despite a loss of topographic organization along the optic nerve (Horton et al., 1979; Colello and Guillery, 1998).

Thus far we have focused on the local retinotopy at the presynaptic input level, and demonstrated a precise topographic organization of the RGC axons in the mouse SC (Figures 2-4). What about the postsynaptic side? Taking a similar approach, we next examined the retinotopy of SC somata at single-cell resolution (Supplemental Figure 3). Specifically, using *in vivo* two-photon calcium imaging, we mapped the RFs of local neurons in the superficial SC layer (58 ± 27 cells/recording from 20 animals; RF size, 5.1 ± 0.9 degrees; RF overlap, 37 ± 11 %; N=1191 cells in total; Supplemental Figure 1B), and performed the same tiling pattern analysis using the Delaunay triangulation (neighboring somata distance, 56 ± 28 µm, N=2292 pairs, Supplemental Figure 1C; neighboring RF distance, 4.1 ± 1.9 degrees, N=2287 pairs, Supplemental Figure 1D). As expected, we found that the tiling patterns of SC somata and their corresponding RFs agreed well in general (77 ± 5%, Figure 4A). However, the agreement was significantly lower for SC somata than for RGC axons (p = 0.001, rank sum test; Figure 4A), indicating that local cellular-level retinotopy is less precise for the postsynaptic neurons than for the input axons. Moreover, the average discrepancies between the observed and retinotopically optimal locations of SC somata (23 ± 5 μm; Figure 4C) or their RFs (1.8 ± 0.3 degrees; Figure 4D) were comparable to those for RGC axons (p = 0.27 and 0.66, respectively; rank sum test), suggesting that the connectivity between RGC axons and SC neurons is not necessarily made to improve the precision of local retinotopy. Thus, our data disagree with the selective connectivity model whereby SC neurons selectively integrate inputs from appropriate presynaptic partners to reconstruct the topography at single-cell resolution (Model 2 in Figure 1). Instead, we suggest that retinotopy in SC arises primarily from precise RGC axonal projections (Model 1 in Figure 1), without much need to elaborate the postsynaptic connectivity.

**Figure 5:**
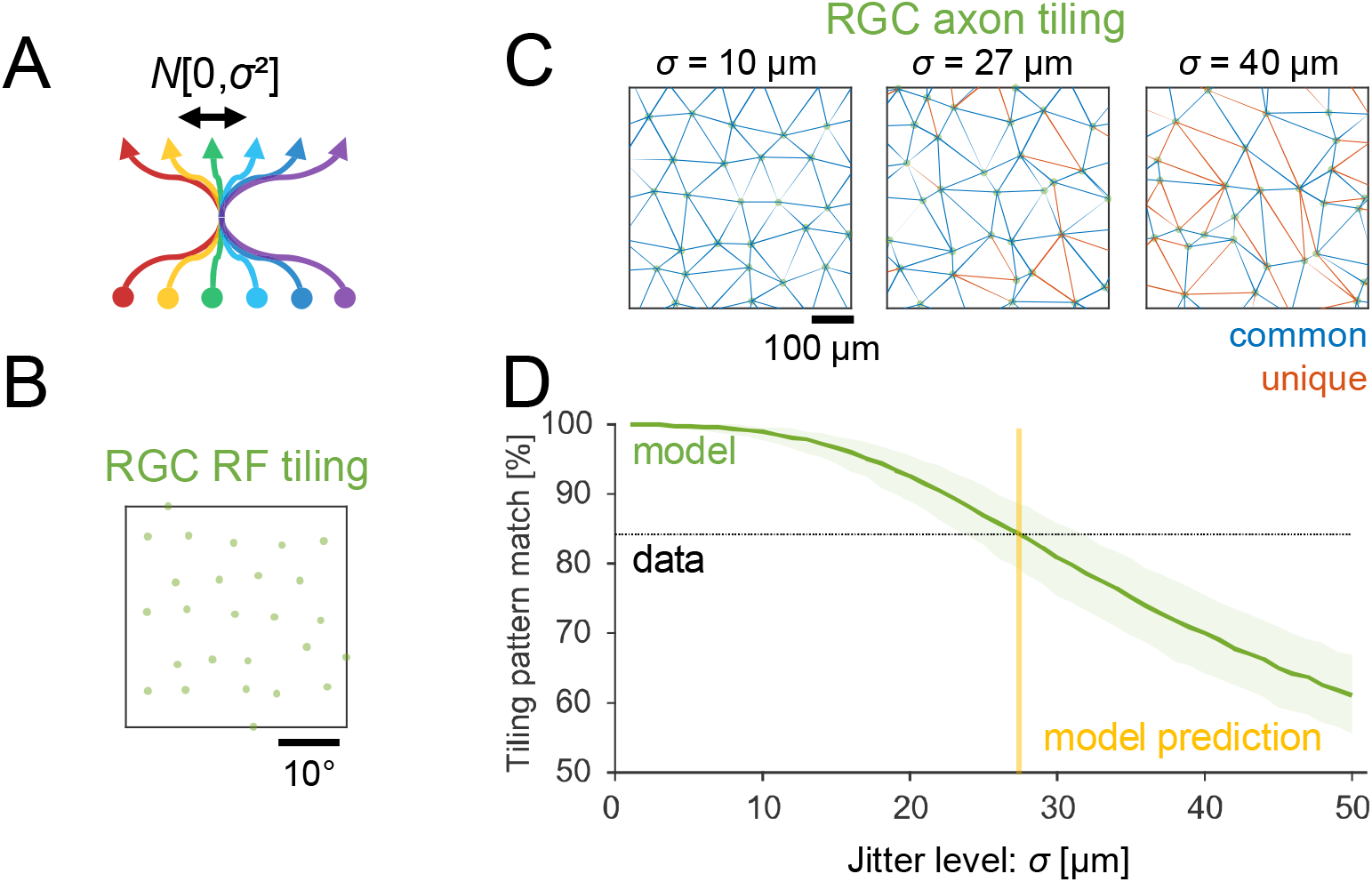
Data-driven model prediction on the precision of retinal ganglion cell axonal projection to the mouse superior colliculus. *A*: Schematic of a retinocollicular projection model. We assumed that 1) a jitter of RGC axonal projection follows a Gaussian distribution *N*[0, σ^2^]; and 2) the tiling pattern of RGC RF centers corresponds to that of the cell locations in the retina. See Methods for details. *B,C*: Representative tiling patterns of simulated RGC RF centers (B; on a 10%-jittered hexagonal lattice) and the corresponding axon centers at different jitter levels (C; *σ* = 10, 27 and 40 μm from left to right panels, respectively). Common (blue) and unique (red) triangulation edges are also shown in each panel of C, when compared to the RF tiling pattern in B. The hexagonal lattice spacing was set to be 7.2 degrees and 100 μm for RGC RFs and axons, respectively, from the experimental data (Supplemental Figure 1). *D*: Correspondence of the triangulation edges between simulated RGC RF and axon tiling patterns at different projection jitter levels (median with 95% confidence interval; 1,000 repetitions). The intersection with the experimentally identified value (horizontal dotted line, 84% from Figure 4A) gives a model prediction on the precision of RGC axonal projection (vertical yellow line; *σ* = 27 ± 4 μm, with 95% confidence interval).

Our tiling pattern analysis showed a near-perfect retinotopy already at the level of the axonal inputs to the mouse SC and no further improvement in the precision of retinotopy for local SC neurons (Figures 3 and 4). What are the conditions to achieve such topographic organizations in the retinocollicular pathway? To address this question, we next performed a computational modelling analysis (see Methods for details). Specifically, by comparing the observed and simulated tiling patterns on the pre- and post-synaptic sides, here we quantified the following two parameters: 1) the precision of RGC axonal projection to a target location in SC (Figure 5); and 2) the number of connecting RGCs to individual SC neurons (Figure 6). Here we did not consider any structural plasticity in our model because the focus is not on the developmental process but on the end result of the axonal organization in adult mice.

We first modelled the tiling patterns of RGC axons at different jitter levels to identify how small the projection error needs to be to recapitulate the observed precision of retinotopy (Figure 5A). The tiling pattern of RGC somata – or equivalently, that of RGC RFs – was simulated as a 2D hexagonal lattice with a small additive Gaussian noise, where the standard deviation of the jitter followed 10% of the lattice spacing to replicate the dense packing of the cell bodies in the retina (e.g., Figure 5B). The tiling pattern of RGC axons in SC was then simulated by introducing additional Gaussian noise to the simulated RGC RF tiling pattern, where the standard deviation σ of this additional noise determines the jitter level of the axonal projection (e.g., Figure 5C). Here we set the axonal lattice spacing to be 100 μm based on our experimental data (Supplemental Figure 1C), and ran the tiling pattern analysis as we did on our experimental data to quantify the precision of retinotopy in the model. As expected, the larger the jitter was, the less precise the retinotopy was (Figure 5D). This allowed us to determine the jitter size that agreed with the observed precision level of retinotopy (84%; Figure 4A): i.e., σ = 27 ± 4 μm (with 95% confidence interval). This is consistent with the average discrepancy between the observed and retinotopically optimal tiling patterns (Δ = 27 μm; Figure 4C), hence validating our modelling framework and further supporting our estimate on the precision of the RGC axonal projection.

**Figure 6:**
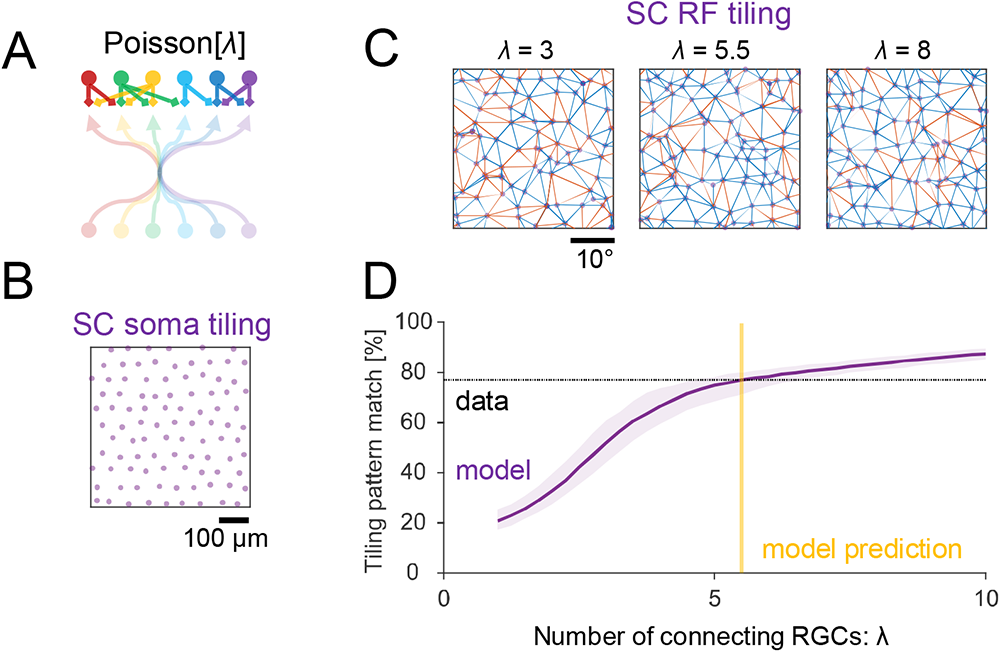
Data-driven model prediction on the number of connecting retinal ganglion cells to individual neurons in the superior colliculus. *A*: Schematic of a retinocollicular mapping model. We assumed that 1) each SC neuron integrates inputs from *λ* nearest neighbor RGC axons, where *λ* follows a Poisson distribution; and 2) the connectivity weights depend on the amount of overlap between SC dendritic field (radius, 200 μm; Gale and Murphy, 2014) and RGC axonic field (135 μm; Supplemental Figure 1A). RGC axonal tiling was simulated with *σ* = 27 μm (Figure 5). See Methods for details. *B,C*: Representative tiling patterns of simulated SC soma centers (B; on a 10%-jittered hexagonal lattice with 56 μm spacing; Supplemental Figure 1C) and the corresponding RF centers at different integration levels (C; *λ* = 3, 5.5, and 8, from left to right panels, respectively), overlaid with common (blue) and unique (red) triangulation edges. *D*: Correspondence of the triangulation edges between simulated SC soma and RF tiling patterns at different input convergence levels (median with 95% confidence interval; 1,000 repetitions). The intersection with the experimentally identified value (horizontal dotted line, 77% from Figure 4A) gives a predicted number of connecting RGCs to individual SC neurons (vertical yellow line; *λ* = 5.5 ± 1.0, with 95% confidence interval).

We next modelled the retinotopy of SC somata on top of the optimal RGC projection model described above (jitter size, σ = 27 μm), using the average number of RGC inputs to SC neurons, λ, as a key model parameter (Figure 6A). The simulated tiling pattern of SC somata (e.g., Figure 6B) was generated in a similar way to that of RGC somata, but with a lattice spacing of 56 μm based on our experimental data (Supplemental Figure 1C). For each simulated SC cell, the RF center location was then determined as a weighted average of the RF centers of neighboring RGC axons (Figure 6C), where we assumed that the number of connecting RGCs followed a Poisson distribution (mean, λ), and that the connectivity strength was proportional to the amount of overlap between the SC cell’s dendritic field (radius, 200 μm; Gale and Murphy, 2014) and the RGC’s axonic field (135 μm; Supplemental Figure 1D). Here we introduced a rather simple connectivity rule as implicated by our experimental data solely from the retinotopy viewpoint (Figures 3 and 4), while details on the cell-type specific connectivity are beyond the scope of our modelling framework. The tiling pattern analysis on the simulated SC cells then showed that the larger the number of connecting RGCs was, the more precise the retinotopy was (Figure 6D). This suggests that the observed relatively less precise retinotopy for SC somata (77% as opposed to 84% for RGC axons; Figure 4A) results from a low input convergence. Indeed, by comparing the model outcome with the experimental data, here we derived that on average SC neurons receive inputs from λ = 5.5 ± 1.0 RGCs (with 95% confidence interval). This is consistent with the observation in the previous electrophysiological studies (Chandrasekaran et al., 2007; Sibille et al., 2022).

## Discussion

Using *in vivo* two-photon axonal imaging, we conducted functional mapping of the retinocollicular projection in adult mice, and demonstrated a precise retinotopic tiling of RGC axon terminals in SC at single-cell resolution (Figures 2 and 3). Here we calculated the projection error size in two different ways: 1) based on the deviation from a linearly-estimated retinotopically-ideal target location (Figures 3 and 4), and 2) by data-driven computational modelling (Figure 5). Both methods consistently found that the projection jitter was below 30 μm (or equivalently 2 degrees of visual angles), much smaller than the observed RGC axonic field size (135 μm; Supplemental Figure 1). These long-range axons can thus be innervated to their exact target locations to faithfully transmit topographic information from the retina, despite a loss of topography in the optic nerve (Horton et al., 1979; Colello and Guillery, 1998). Our results highlight the precision of the developmental processes, from genetically-determined sorting of RGC axons (Plas et al., 2005; Dhande et al., 2011) to activity-dependent structural plasticity in SC (Chandrasekaran et al., 2005).

In contrast, we found that the local retinotopy of SC somata was no better than that of RGC axons (Figure 4 and Supplemental Figure 3). The connectivity between RGCs and SC cells is thus not necessarily made to retain or improve the topography. Instead, assuming no selectivity in the connectivity patterns, our modelling analysis indicates that a reduced precision of local retinotopy on the postsynaptic side can be a direct consequence of a low input convergence level (Figure 6). Based on our experimental data, we derived from our model that on average SC neurons receive inputs from ∼5.5 RGCs (Figure 6). While connectivity patterns between specific RGC and SC cell types remain an open question, this is consistent with the past electrophysiological measurements (Chandrasekaran et al., 2007; Sibille et al., 2022), justifying our model framework and conclusion.

Taken together, we suggest that retinotopy in the mouse SC arises largely from topographically precise projection of RGC axons, rather than local circuit computation by SC neurons. While here we studied only the medial-posterior part of the superficial SC (Figure 2), we expect that this type of organization exists across SC, given that the mechanisms underlying the retinotopy development are not dependent on the spatial location of SC (Cang and Feldheim, 2013; Arroyo and Feller, 2016). Note, however, that the precision of axonal projection was not perfect. Postsynaptic circuit mechanisms should then be indispensable as well in retaining topography, especially for higher-order processing because otherwise retinotopy will no longer be recognizable after a cascade of signal transmission along the visual hierarchy (e.g., below chance level after six ∼80% precision transmissions). It is a future challenge to investigate the cellular-level topographic organization in other brain areas, including the retinotopy in the downstream visual pathways (Wandell et al., 2007), and clarify the contribution of pre- and post-synaptic circuit mechanisms in each area.

Having a precise retinotopy at single-cell resolution facilitates spatial information processing not only at a global level, but also at local circuit levels. For example, looming detection has been suggested to arise de novo in the superficial SC layer (Lee et al., 2020). In principle, this can be achieved even in the absence of retinotopy by elaborating the wiring among local neurons. It is, however, much more efficient to exploit precise topographic information conveyed from the retina because the connectivity length and its complexity can be minimized to locally process spatial information at any point in the visual field. This will also help align different topographic maps in the same brain area to function coherently, such as the retinotopy, orientation, and ocular dominance maps in SC (Feinberg and Meister, 2015; de Malmazet et al., 2018). The observed precise spatial organization we demonstrate here suggests that the wiring efficiency indeed matters for local circuit computation.

How can then such a precise retinotopic projection be formed? Retinotopic map formation in SC occurs during the first postnatal week in mice, involving both genetic and activity-dependent factors (Cang and Feldheim, 2013; Arroyo and Feller, 2016). These factors also play a key role in the development of a fine-scale organization in other sensory systems, such as the tonotopic map in the cochlear nucleus (Krasewicz and Yu, 2023), and the chemotopic map in the olfactory bulb where olfactory sensory neurons expressing the same olfactory receptor type project exclusively to the same single glomerulus (Imai et al., 2010). While overall sensory map formation is genetically predetermined by molecular cues (e.g., ephrin-Eph signaling; Frisén et al., 1998; Krasewicz and Yu, 2023; and axon-axon interactions; Plas et al., 2005; Cioni et al., 2018), a precise topography is established only after refinement that involves spontaneous activity, such as retinal waves during development (Chandrasekaran et al., 2005; Arroyo and Feller, 2016), and eventually experience-driven alignment (McLaughlin et al., 2003). In particular, here we suggest that this refinement process of the retinocollicular projection during development should be extremely precise, to the extent that retinotopy arises at a single-cell resolution in adult animals (Figures 3 and 4). It is then possible that neuronal circuits are in general wired more precisely than previously thought to exploit topographic information for their function, including long-range projections to other retinal targets (Liang, et al., 2018) as well as those in other systems, such as the callosal projections and entorhinal-hippocampal networks (Fame et al., 2011; Jbabdi et al., 2013; Patel et al., 2014).

## Methods

No statistical method was used to predetermine the sample size. All experiments involving animals were performed under the license 233/2017-PR from the Italian Ministry of Health, following protocols approved by the Institutional Animal Care and Use Committee at European Molecular Biology Laboratory. The data analyses were done in Python and Matlab (Mathworks). The statistical significance level was set to be 0.05. All summary statistics were described as median ± median absolute deviation unless otherwise noted.

### Animals

Female C57BL/6J mice (*Mus musculus*; 15 for axonal imaging; 20 for somatic imaging) were used at around 6-10 weeks of age at the time of the first surgery. Mice were kept on a 12-h light / 12-h dark cycle and given water and food *ad libitum*. After surgery, the animals were kept in groups operated on the same day. Mice were between 12-24 weeks of age at the time of imaging experiments.

### Intravitreal viral injections

Intravitreal injection of recombinant adeno-associated virus (AAV) 2, pseudotyped with a hybrid of AAV1 and AAV2 capsids, was used to deliver hSyn-axon-GCaMP6s expression cassette to the mouse retinal ganglion cells (RGCs; Broussard et al., 2018). For the viral injection, mice were anaesthetized (induction, 4% isoflurane in oxygen; maintenance, 1.8-2.0%) and kept on a heated plate (Supertech Physiological Temperature Controller) to avoid hypothermia. Both eyes were protected by saline drop or viscous eye ointment (VitA-POS, Ursapharm). The scleral surface on the left eye was exposed and a small piercing was made with a sterile 28-30G needle in between the sclera and the cornea. An injection pipette (∼50 µm tip diameter with 30-40° bevel) prefilled with a virus solution (∼1.5×10^14^ vg/mL in phosphate-buffered saline with 0.001% Pluronic F68 and 0.001% FastGreen) was then inserted into the vitreous chamber approximately 1 mm deep. The injection pipette was made from a borosilicate glass capillary (1B120F-3, WPI) with a pipette puller (DMZ, Zeitz) and a microgrinder (EG-45, Narishige). After a good sealing of the pipette was formed, 1.2 µL of the virus solution was injected at a rate of 10 nL/s using a microinjection pump (either Neurostar NanoW or WPI NanoLiter 2010) with mineral oil (Sigma, M5904) filled in the displacement space by a stainless steel plunger. The pipette was slowly withdrawn at least 5 minutes after the completion of the injection, and the treated eye was covered with the eye ointment. The animal was then allowed to recover from anesthesia in a warmed-up chamber and brought back to its home cage.

### Intracranial viral injections

Pseudotyped AAV, composed of AAV2 rep and AAV9 cap genes or a hybrid of AAV1 and AAV2 cap genes, was locally injected to the mouse superior colliculus (SC) for the expression of genetically-encoded calcium indicators (jGCaMP7f, jGCaMP8m or jRGECO1a) under pan-neuronal human synapsin (hSyn) promoter. The intracranial viral injection was made at the same time as the cranial implantation as described below. After making a craniotomy over the right SC, an injection pipette (∼30 µm tip diameter; WPI 1B120F-3 borosilicate glass capillary pulled with Zeitz DMZ puller) prefilled with a virus solution (∼5×10^12^ to ∼4×10^14^ vg/mL in phosphate-buffered saline) was inserted across the dura at coordinates from Bregma around -4 mm AP, 0.5-0.7 mm ML, and then slowly advanced until ∼1.25 mm deep. The virus solution (0.4-0.6 µL) was injected at a rate of 2 nL/s with a microinjection pump (either Neurostar NanoW or WPI NanoLiter 2010). The pipette was slowly withdrawn at least 10 minutes after the completion of the injection, followed by the cranial window implantation procedure.

### Cranial implantations

We adapted methods described in Feinberg and Meister (2015) for the cranial window implantation over the mouse SC. A cranial window assembly was made in advance, where the surface of a circular glass coverslip (5 mm diameter, 0.13-0.15 mm thickness; Assistent Karl Hecht) was activated by a laboratory corona treater (BD-20ACV Electro-Technic Products) and fused to a cylindrical silicone plug (1.5 mm diameter, 0.75 – 1.00 mm height; Kwik-Sil, WPI) by baking it for 24 hours at 70-80°C.

For the implantation, animals were anaesthetized (induction, 4% isoflurane in oxygen; maintenance, 1.5-2.0%) and placed inside a stereotaxic apparatus (Stoelting 51625). Throughout the surgery, temperature was maintained at 37°C using a heated plate (Supertech Physiological Temperature Controller) to avoid hypothermia, and the eyes were protected with eye ointment (VitA-POS, Ursapharm). After disinfecting and removing the scalp (Betadine 10%, Meda Pharma), the skull surface was scratched and cleaned to ensure good cement adhesion. A craniotomy of a size about 3.0 mm (anterior-posterior; AP) by 2.5 mm (medial-lateral; ML) was made over the right SC using a high-speed surgical drill (OmniDrill35, WPI) with a 0.4 mm ball-tip carbide bur (Meisinger). To prevent bleeding, the craniotomy was treated by hemostatic sponges (Cutanplast, Mascia Brunelli) soaked with sterile cortex buffer (NaCl 125 mM, KCl 5 mM, Glucose 10 mM, HEPES 10 mM, CaCl2 2 mM, MgSO4 2 mM, pH 7.4).

For SC somata imaging, viral injections were made as described above. The implant was then placed in a way to push the transversal sinus and posterior cortex ∼0.5 mm forward and position the silicone plug over the medial-caudal region of the right SC. Tissue adhesive (Vetbond, 3M) was used to fix and seal the implant. A custom-made titanium headplate (0.8 mm thick) was then cemented to the skull using acrylic cement powder (Paladur, Kulzer) pre-mixed with cyanoacrylate adhesive (Loctite 401, Henkel).

After the surgery, the animal was recovered from anesthesia in a warmed-up chamber and returned to its home cage. For postoperative care, animals were given intraperitoneally 5 mg/kg Rimadyl (Zoetis) and 5 mg/kg Baytril (Bayer) daily for 3-5 days. We waited for another 10-15 days until the cranial window completely recovered before starting *in vivo* two-photon imaging sessions (e.g., Figure 2B).

### Visual stimulation

Visual stimuli were presented to the subject animals as described previously (Boissonnet et al., 2022). In short, a custom gamma-corrected digital light processing device was used to project images (1280-by-720 pixels; frame rate, 60 Hz) to a spherical screen (radius, 20 cm) placed ∼20 cm to the contralateral side of an animal’s eye, stimulating the visual field ±22° in elevation and ±36.5° in azimuth (Figure 2A and Supplemental Figures 2A and 3A). We presented 1) random water-wave stimuli (2-10 min) for generating binary masks for signal source extraction in calcium image analysis (see below); and 2) randomly flickering black-and-white checkerboard stimuli (10 min) for receptive field mapping, with rectangular fields 3.7° in width and 2.9° in height, each modulated independently by white noise at 4 Hz. When these stimuli were not presented, the screen remained uniformly grey to keep the average light intensity level constant.

### *In vivo* two-photon imaging

Prior to *in vivo* imaging sessions, animals were habituated to stay head-fixed on a custom-made treadmill disc (8-10 habituation sessions in total over a week, each for 2 hours). For the imaging session, animals were kept on the treadmill with their head fixed for no longer than 2 hours (2-5 sessions/animal). Two-photon calcium imaging was done on a galvo-resonant multiphoton microscope (Scientifica HyperScope with SciScan image acquisition software) equipped with a mode-locked tunable laser (InSight DS+, Spectra-Physics) and a plan fluorite objective (CFI75 LWD 16X W, Nikon). In each imaging session, we performed single-plane time-lapse recordings (field of view, approximately 0.65-by-0.65 mm) at a depth of 120-220 μm from the SC surface. The fluorescent signal (excitation wavelength, 920 nm for axon-GCaMP6, GCaMP7f, and GCaMP8m; 1040 nm for jRGECO1a; average laser power under the objective, 40-80 mW) was bandpass-filtered (BP 527/70 or BP 650/100 after beam-splitter FF580-FDi01, Semrock) and detected with a non-descanned gallium arsenide phosphide photomultiplier tube (Hamamatsu GaAsP PMT). Each frame was acquired with 1024-by-1024 pixels (16-bit depth) at 15.4 Hz for RGC axonal imaging, and 512-by-512 pixels (16-bit depth) at 30.9 Hz for SC somata imaging.

### Calcium image analysis

For preprocessing of RGC axon data, the original 1024-by-1024 pixel images were first downsampled to 512-by-512 pixels (2-by-2 pixels averaging) to reduce noise. To correct motion artefacts, we performed two iterations of Fourier-based rigid image registration in ImageJ, followed by cropping the image border by 16 pixels; and then ten iterations of non-rigid motion correction (NoRMCorre) in CaImAn (Giovannucci et al., 2019), followed by a 12-pixel border crop. The resulting images (456-by-456 pixels; e.g., Supplemental Movie 1) represent a field of view of around 0.57-by-0.57 mm (1.3 μm/pixel).

From the preprocessed images, we identified the axonal patches of individual RGCs and extracted their signals in CaImAn (e.g., Figure 2C,D). Specifically, using a part of the recordings (3,000-5,000 frames representing the random water-wave stimulus presentation period), we first ran two iterations of constrained non-negative matrix factorization (CNMF) in CaImAn, where we set the number of expected components (*params.K*) to be 60 as an initialization parameter. From the identified components, we then manually selected those with a uniformly-filled oval-like shape that had a size of around 50-150 μm as biologically relevant ones (Hong et al. 2011), and converted them into binary spatial masks to run two iterations of masked CNMF for processing the entire time-lapse recordings. The resulting set of spatial components (*estimates.A*) and deconvolved neural activities (*estimates.S*) was used for the subsequent analyses.

The area of the individual RGC axonal patches *P_i_* was estimated from the identified spatial components in CaImAn (1.3 µm/pixel), from which the radius was estimated as (*P_i_*⁄π)^0.5^ under the assumption of a circular patch shape (Supplemental Figure 1A). The fraction of the overlap between identified axonal patches was calculated as the ratio of the areas between the intersection of any two patches *U_i≠j_*(*P_i_* ∩ *P_j_*) and the union of all patches ⋃_*i*_ *P_i_*.

For SC soma data, we first ran a sequence of the rigid and non-rigid motion corrections in CaImAn, followed by image cropping from 512-by-512 pixels into 480-by-480 pixels in ImageJ (1.3 μm/pixel). Using a part of the recordings (3,000 frames from the random water-wave stimulus presentation period), we then ran two iterations of CNMF in CaImAn, where the images were divided into 6-by-6 (36 in total) patches and the expected number (*params.K*) and size (*params.gSig*) of neurons were set to be 5 per patch and 5-by-5 pixels in half size, respectively, as initialization parameters. From the identified putative cells, we manually selected those with a uniformly-filled round shape of around 10-20 μm in size as biologically relevant ones, and converted them into binary spatial masks to run two iterations of masked CNMF in CaImAn on the entire time-lapse recordings. The resulting set of spatial components (*estimates.A*) and deconvolved neural activities (*estimates.S*) was used for the subsequent analyses.

### Receptive field analysis

The receptive fields (RFs) of the identified RGC axon patches or SC somata were estimated by reverse-correlation methods using the random checkerboard stimuli (Chichilnisky, 2001). Specifically, we calculated the response-weighted average of the stimulus waveform (0.5 s window; 1/60 s bin width), and characterized its spatial profile by the two-dimensional (2D) Gaussian curve fit at the peak latency (e.g., Figure 2E and Supplemental Figures 2B and 3B). The RF center was assigned to the center of that 2D Gaussian profile, and the RF size was estimated as twice the mean standard deviation (SD) of the long and short axes (Supplemental Figure 1B). The fraction of the overlap between the RFs (1 SD Gaussian profiles) was computed similarly as for the axonal patches. The Pearson correlation coefficient between the RF size and the RGC axonal patch size was calculated with the 95% interval of the data to eliminate the outliers. Those cells that had the RF center on the border or outside the stimulus screen were eliminated from the tiling pattern analysis described below. Those recordings that had less than 10 cells with RF centers on the stimulus screen were also excluded from the tiling pattern analysis.

### Tiling data analysis

To compare the tiling patterns between RGC axon patches / SC somata and their RFs (e.g., Figure 3 and Supplemental Figures 2 and 3), we first computed the Delaunay triangulation of their centroid locations using the Euclidean distance in each space (e.g., Figure 3C,D and Supplemental Figures 2C,D and 3C,D). As a measure of similarity between the two tiling patterns, we then calculated the number of common edges, divided by the mean of the total number of edges in each triangulation. This measure is referred to as “tiling pattern match” in Figures 4-6. The chance level was calculated by a bootstrap method (10,000 repetitions; Figure 4A).

We used the least squares method to identify an optimal Affine transformation for mapping a given RF tiling pattern onto the corresponding tiling pattern of RGC axons (e.g., Figure 3E), or vice versa (e.g., Figure 3F). The Euclidean distance of the cell or RF locations between the observed and affine-transformed tiling patterns was then used as a measure of the precision of local retinotopy (Figure 4C,D).

### Modelling of retinocollicular mapping

We modelled the retinocollicular mapping in four steps (Figures 5 and 6).

1. The tiling pattern of RGC somata was simulated as a 2D hexagonal lattice with a Gaussian jitter σ [0, σ^2^]. The standard deviation *σ* was set to be 10% of the lattice spacing *L* to recapitulate the dense packing of the cell bodies in the retina. We assumed that the tiling pattern of RGC RFs was equivalent to the corresponding somatic tiling pattern (e.g., Figure 5B; *L* = 7.2 degrees from Supplemental Figure 1D).
2. The tiling pattern of RGC axons in SC (i.e., retinocollicular projection) was then simulated by introducing additional Gaussian noise *N*[0, σ^2^] to the RGC RF tiling pattern from step 1 (e.g., Figure 5C) but with *L* = 100 μm (Supplemental Figure 1C). When *σ* = 0 μm, the tiling pattern of RGC axons is identical to the somatic tiling pattern, showing perfect retinotopy (i.e., 100% tiling pattern match).
3. The tiling pattern of SC somata was simulated as a 2D hexagonal lattice (*L* = 56 μm; Supplemental Figure 1C) with a Gaussian jitter (*σ* = 0.1*L*) as in step 1 (e.g., Figure 6B).
4. The RF of each SC neuron was calculated by integrating inputs from *λ* nearest neighbor RGC axons, where *λ* follows a Poisson distribution. Specifically, assuming that the connectivity strength depends on the amount of overlap between SC dendritic field (radius, 200 μm; Gale and Murphy, 2014) and RGC axonic field (135 μm; Supplemental Figure 1A), we defined the SC RF center location as the weighted average of the RF centers of the connecting RGCs (e.g., Figure 6C).

To identify the precision of RGC axonal projection to SC (Figure 5), we ran the steps 1 and 2 at different jitter levels *σ* (from 0 to 50 μm in steps of 1 μm; 1,000 repetitions each) and calculated the similarity between the simulated tiling patterns of RGC axons and their RFs using the triangulation method as described above (Figure 5D). We then determined the jitter level *σ* where the simulated tiling pattern similarity agreed with the experimental data (84%; Figure 4A).

To estimate the average number of connecting RGCs to individual SC neurons (Figure 6), we ran the steps 1-4 at different mean values *λ* (from 1 to 10 RGCs in steps of 0.25; 1,000 repetitions; *σ* = 27 μm for step 2 from Figure 5D) and calculated the similarity between the simulated SC somatic and RF tiling patterns using the triangulation method as described above (Figure 6D). We then determined the input convergence level *λ* where the simulated tiling pattern similarity agreed with the experimental data (77%; Figure 4A).

## Acronyms

2D: two-dimensional
AAV: adeno-associated virus
CNMF: constrained non-negative matrix factorization
RF: receptive field
RGC: retinal ganglion cells
SC: superior colliculus
SD: standard deviation

## Acknowledgments

This work was supported by research grants from EMBL (H.A.). The EMBL Genetic and Viral Engineering Facility is acknowledged for virus production; EMBL IT Support for provision of computer and data storage servers; and the LAR facility for taking care of animals. We thank Lin Tian (University of California Davis) for providing axon-targeted GCaMP construct; Thomas Euler and Luke Rogerson (University of Tübingen) for sharing Python scripts to generate random water-wave stimuli, originally from Andreas Tolias lab (Baylor College of Medicine); Simone Calabrese, Ilaria Sauve, Grace Cunliffe, and Matteo Tripodi for supporting experiments; and all the Asari lab members as well as Santiago Rompani for many useful discussions.

## Author contributions

D.M. and H.A. designed the study; D.M. and L.F performed experiments; D.M, T.B., L.F. and H.A. analyzed the results; and D.M., L.F. and H.A. wrote the manuscript.

